# BeeMonitor: Automated IoT video surveillance and an AI-powered video processing system for monitoring the foraging and nesting behavior of cavity-nesting solitary bees

**DOI:** 10.64898/2026.07.10.737879

**Authors:** Edward Idun Amoah, Santosh Sanjel, Natalie Boyle, Christina Grozinger

## Abstract

1. Solitary bee species that use artificial trap nests are important for agricultural crop production and as indicators of habitat quality. Quantifying cavity-nesting solitary bee foraging and nesting behavior is essential for real-time analysis of population numbers and pollination activity, as well as understanding how environmental conditions shape reproductive success and population dynamics. However, manual observation is labor-intensive, prone to observer bias, and unable to deliver continuous data. Existing automated systems either require individual bee marking or detect presence without resolving nest-tube-level entry and exit events.
2. We developed BeeMonitor, an integrated hardware and computer-vision pipeline that detects nest entry and exit events in cavity-nesting solitary bees from continuous video, using *Osmia cornifrons* (the horn-faced mason bee) as a model system. A low-cost Raspberry Pi handles solar-powered field recording, while the software combines object detection (YOLOv26), a custom multiple-object tracker (BeeTrack), and a Random Forest classifier trained on trajectory-derived features to distinguish genuine events from incidental detections.
3. Over a 29-day deployment, hardware reliability averaged 97.5% recording coverage. The pipeline achieved 91.3% precision and 87.3% recall (F1 = 0.893), generalizing robustly under leave-one-video-out cross-validation (mean F1 = 0.904). Detected foraging trips correlated strongly with brood cell counts (R2 = 0.849, *p* < 0.001, *n* = 19), and a Random Forest model (AUC = 0.820) identified solar radiation as the dominant driver of foraging activity, followed by temperature.
4. BeeMonitor demonstrates that automated computer vision can reliably extract ecologically relevant behavioral data from continuous video, enabling real-time analysis of pollinator behavior and abundance at a temporal and spatial resolution unattainable through manual observation. Its modular design supports adaptation to other species and monitoring contexts.

## 1. Introduction

Pollinators play a critical role in both natural ecosystems and agricultural production, contributing to the reproduction of approximately 80% of flowering plant species and the production of about 35% of global crops (Eilers et al., 2011; Jordan et al., 2021; Rodger et al., 2021). Despite their ecological and economic importance, many pollinator populations are experiencing widespread declines driven by multiple interacting stressors, including habitat loss, pesticide exposure, climate variability, and reductions in floral resources (Cornelisse et al., 2025; Dicks et al., 2021; Wagner et al., 2021). Understanding how environmental factors affect pollinators is critical for advancing ecological research, as these drivers shape behavior, physiology, survival, and ultimately reproductive success and population dynamics (Gérard et al., 2024; Henry et al., 2012; Kenna et al., 2021; McKinney & Park, 2012; Siviter et al., 2021). These responses, however, are not uniform; different pollinator species vary considerably in their sensitivity to environmental change, owing to differences in life history traits, physiology, and phenology (Hamblin et al., 2017; Slominski & Burkle, 2019; Williams et al., 2010), making species-specific monitoring essential for predicting how pollinator communities will be affected under different environmental scenarios.

Solitary bee species that nest in above-ground cavities serve both as important pollinators of agricultural crops and indicators of habitat quality (Osterman et al., 2021; Staab et al., 2018). Indeed, an increasing number of studies use artificial trap nests with hollow tubes to evaluate bee abundance and species diversity in response to anthropogenic and environmental change (Staab et al., 2018). For cavity-nesting solitary bees such as *Osmia cornifrons*, reproductive success depends on the construction and provisioning of brood cells within nests (Bosch & Kemp, 2001; Ganser et al., 2021). Each brood cell requires multiple foraging trips to first collect the construction materials for the cell (in the case of *O. cornifrons*, mud) and additional trips to collect pollen and nectar, upon which the egg is deposited (Bosch et al., 2008; Klostermeyer & Gerber, 1969). Despite the importance of foraging behavior as a proxy for reproductive dynamics, quantifying this behavior in natural systems remains challenging. Most field studies rely on manual observations conducted over short time windows and limited spatial extent or coverage (Bonelli et al., 2020; Bosch, 1994; Goodell, 2003; McKinney & Park, 2012). These approaches are labor-intensive and limit spatiotemporal resolution, constraining the ability to capture species-specific variability in activity across environmental conditions and locations. Fine-scale behavioral responses to local microclimate and landscape context are therefore likely to be missed or underestimated.

Advances in Internet of Things (IoT) devices and computer vision offer new opportunities to automate the monitoring of foraging and nesting activity in natural habitats (Nisal Ratnayake et al., 2021). Recent studies have applied computer vision to track bee movement and activity in cavity-nesting environments, such as bee hotels, thereby enabling detection of nest entry and exit events corresponding to individual foraging trips (Knauer et al., 2022). However, existing systems either require an individual bee tag or apply object detection to untagged populations with limited accuracy, without extracting discrete nest-tube-level activity (Knauer et al., 2022; Rozenbaum et al., 2024; Wittmann et al., 2024). Additionally, most of these systems focus on software for a specific use case and are often presented without a thorough evaluation of their field hardware, making it difficult for other ecologists to adopt and adapt them across different contexts. Therefore, there is a need for an integrated modular hardware-and-software system that can be thoroughly tested and adapted to different use cases for field monitoring of bee behavior.

Here, we present BeeMonitor, an integrated hardware-and-software system for automated monitoring of nest-tube-level foraging activity in cavity-nesting solitary bees, without requiring individual marking. By combining off-the-shelf hardware systems like Raspberry Pis with state-of-the-art object detection, custom multi-object tracking, and machine learning–based event classification, BeeMonitor enables continuous, non-invasive quantification of foraging activity from monitoring video data. More broadly, this study introduces a modular generalizable framework for extracting discrete behavioral events from continuous observations, linking individual activity to ecological processes at fine spatiotemporal scales. This study has five objectives: (1) design and build a low-cost IoT hardware system for continuous field monitoring, (2) develop a computer vision software pipeline to extract foraging activity from video data, (3) deploy BeeMonitor to monitor nesting bees in the field, (4) evaluate software pipeline performance against manually annotated ground truth data, and (5) quantify the influence of different environmental conditions on the foraging behaviour of *Osmia cornifrons* with machine learning.

## 2. Materials and Methods

### 2.1 Hardware system

The BeeMonitor hardware system was designed for continuous, field-deployable video monitoring of cavity-nesting bees. The system comprises two modular components: (1) a video recording module and (2) an energy supply module (Figure 1A). This modular design allows flexible deployment under both grid-powered and off-grid conditions.

**Figure 1.**
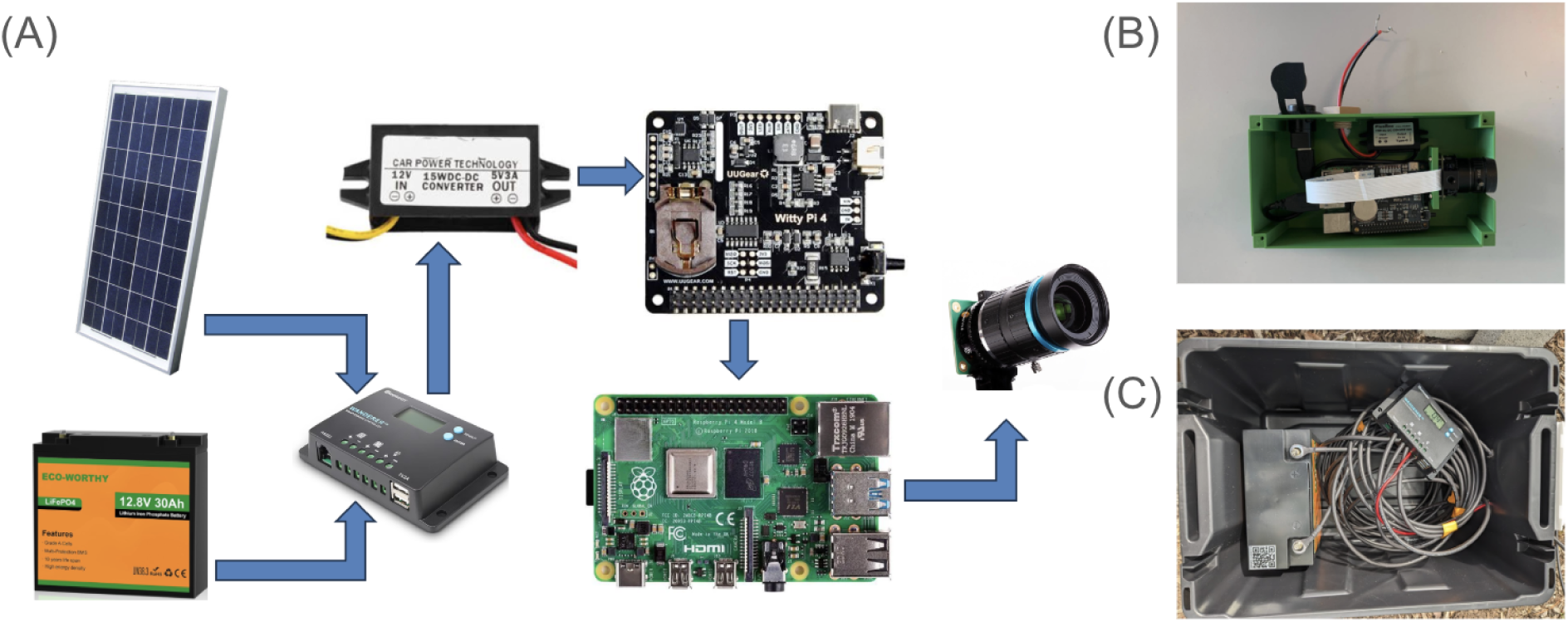
BeeMonitor hardware system. (A) System schematic showing the two modular components: the video recording module (Raspberry Pi 4, IMX477 camera, Witty Pi 4 power management, and SD card storage) and the energy supply module (100W solar panel, charge controller, DC-DC converter, and lithium battery). (B) Assembled video recording modules housed in a weatherproof enclosure. (C) Complete off-grid deployment energy modules integrated in a field-ready enclosure.

The video recording module is built around a Raspberry Pi 4 single-board computer equipped with a high-resolution camera (IMX477 sensor) and programmable power management (Witty Pi 4), enabling automated daily recording cycles (Figure 1B). Video is captured at 30 frames per second and stored in 10-minute segments to balance temporal resolution with storage efficiency. The recording software, implemented in Python, runs as a background service and initiates automatically on system startup.

The energy module consists of a solar-powered system including a photovoltaic panel, charge controller, and lithium battery, providing sufficient capacity for continuous operation in remote field environments (Figure 1C). Detailed hardware specifications, assembly instructions, and software setup are provided on the project GitHub repository.

### 2.2 Software pipeline

The BeeMonitor software pipeline converts continuous video recordings into discrete behavioral events through three sequential modules: object detection, multiple-object tracking, and activity extraction (Figure 2). Each module is implemented as an independent module, allowing individual components to be replaced or reconfigured for different species, camera setups, or ecological contexts without modifying the rest of the pipeline.

**Figure 2.**
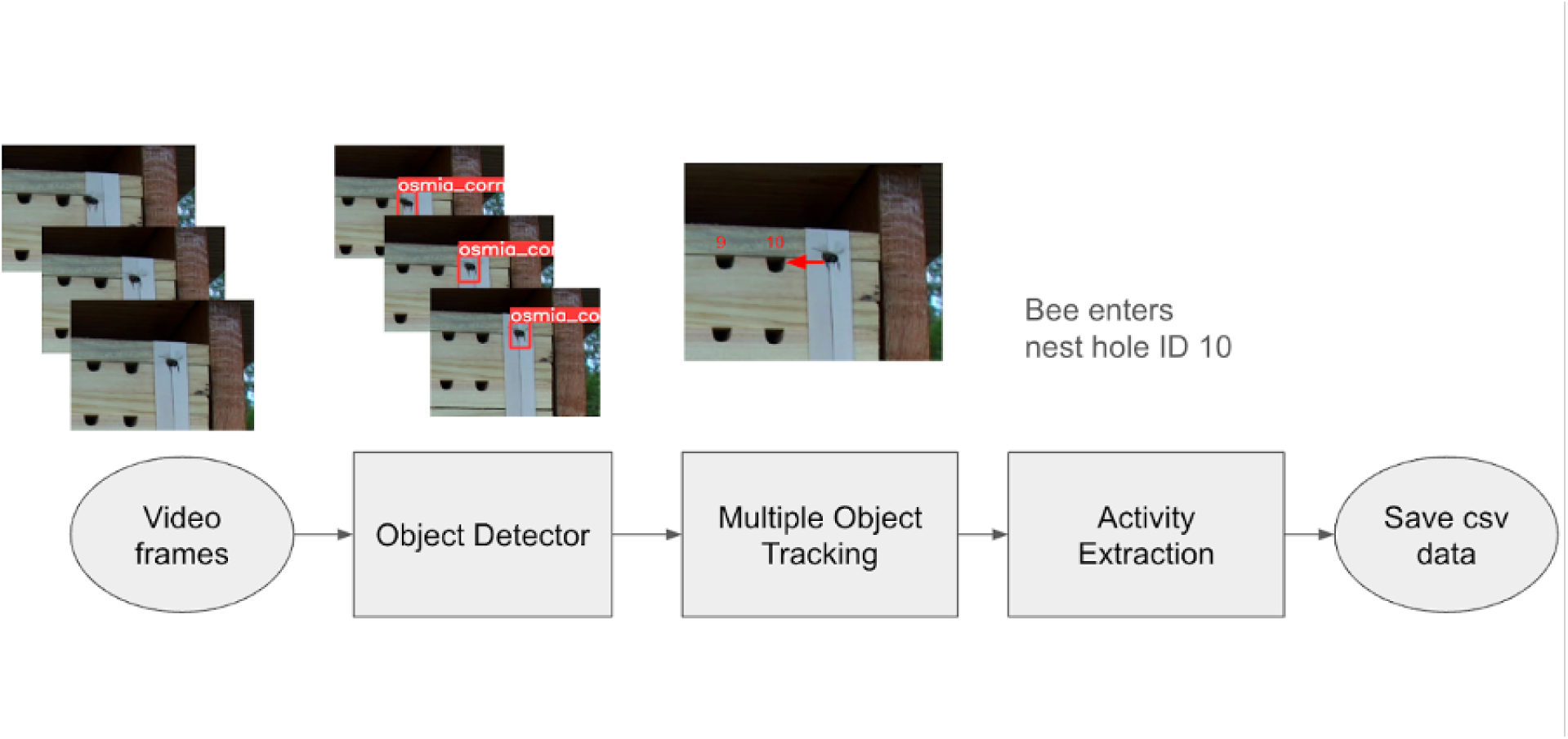
BeeMonitor software pipeline. Video frames are processed sequentially through three modules: object detection (localizing bees and nest tubes in each frame using YOLOv26), multiple-object tracking (reconstructing individual bee trajectories across frames using BeeTrack, a custom SORT-based algorithm with adaptive motion uncertainty), and activity extraction (identifying nest entry and exit events from trajectories using geometric assignment and machine learning classification). Detected events are exported as structured CSV data for downstream ecological analysis. The example shows a bee entering nest hole ID 10.

In the first module, each video frame is processed by a deep learning–based object detector trained to detect nest tubes and bees. We used YOLOv26 (Sapkota & Karkee, 2026), a state-of-the-art single-stage object detection architecture that performs end-to-end inference without requiring post-processing steps such as non-maximum suppression, reducing detection latency and simplifying deployment on resource-constrained hardware. Bee detections provide the raw positional observations used for tracking, while nest tube detections establish the spatial reference frame for event assignment. The detection model can be retrained for different bee species, nest structures, or monitoring contexts, making this module independently adaptable across ecological systems.

In the second module, per-frame bee detections are linked across time using BeeTrack, a custom multiple-object tracking algorithm built on the SORT framework (Bewley et al., 2016). The algorithm operates in two sequential steps per frame (i.e., prediction and association). In the prediction step, a Kalman filter propagates each active track forward using a constant-velocity motion model, estimating where each bee is expected to appear in the next frame based on its current position and velocity (Kalman, 1960). In the association step, the Hungarian algorithm finds the globally optimal assignment between predicted track positions and incoming detections by minimizing a combined cost function that incorporates both spatial distance and bounding-box overlap (Kuhn, 1955).

Standard SORT assumes constant velocity between frames, an assumption suited to pedestrians and vehicles but violated by foraging bees, which frequently change direction abruptly and traverse distances between frames that far exceed their body size (Bewley et al., 2016; Cao et al., 2023; Wojke et al., 2017), leading to poor state parameter updates and fragmented trajectories (Nisal Ratnayake et al., 2021; Rozenbaum et al., 2024). Under standard settings, the Kalman filter motion model places high confidence in its velocity predictions and resists correction by detections that deviate significantly from the expected position, leading to identity switches or track loss during rapid directional changes.

BeeTrack addresses this by increasing the uncertainty of motion predictions, causing the tracker to weigh incoming detections more heavily than predicted positions during each update. This allows tracks to follow abrupt directional changes without identity loss. The matching distance threshold is also expressed as a multiple of the automatically estimated bee size, rather than a fixed pixel value, thereby scaling adaptively across different camera resolutions and deployments. These adaptive features are intentionally implemented to make the software adaptable to different use cases and contexts. Bee size is estimated continuously from incoming detections using a robust outlier-rejection procedure. Dead tracks are briefly retained to allow resurrection if a matching detection reappears within a defined spatial and temporal window, thereby maintaining identity continuity through brief occlusions or missed detections. The tracker can be configured for different movement characteristics or replaced with alternative tracking algorithms as needed.

In the final module, reconstructed trajectories are analyzed to identify foraging events through a two-step process. First, candidate events are identified geometrically. A trajectory whose endpoint falls within the spatial boundary of a nest tube is labeled a candidate entry foraging event, while a trajectory whose starting point falls within a nest tube boundary is labeled a candidate exit foraging event. This step produces a pool of candidates that includes both genuine foraging events and noise trajectories arising from partial occlusions and trajectory fragmentations. Second, a machine learning classifier evaluates each candidate trajectory and assigns a confidence score reflecting the likelihood that it represents a genuine foraging event. The classifier operates on 20 trajectory-derived features organized into five categories: trajectory geometry, speed dynamics, nest proximity, positional variance, and movement direction. Candidates exceeding a confidence threshold are retained as detected events; low-confidence candidates are discarded. Detected events, including event type, nest tube identifier, and timestamp, are exported as structured CSV data for downstream analysis. The event detection logic can be reconfigured to define context-appropriate behavioral events beyond nest entry and exit, supporting adaptation to other monitoring scenarios such as flower visitation or hive entrance activity (Ratnayake et al., 2021; Rozenbaum et al., 2024).

### 2.3 Field deployment evaluation

To evaluate the BeeMonitor system under real-world conditions, we deployed the hardware at a bee hotel at the Penn State Horticulture Research Farm (Mendel’s Way Farm & Greenhouses; 40.8106° N, −77.8600° W, Pennsylvania, USA) during the 2024 nesting season (Figure 3A).

**Figure 3.**
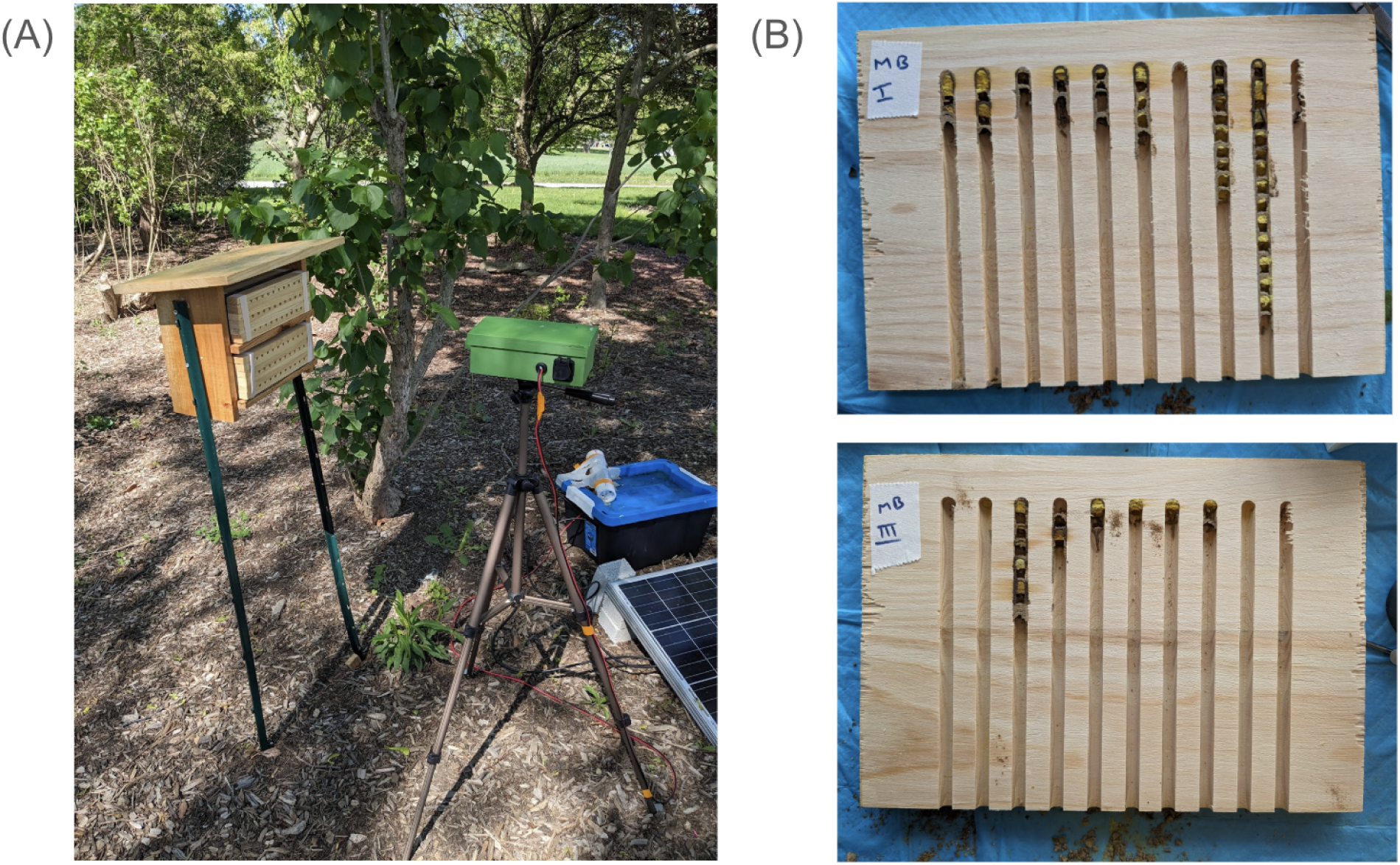
Field deployment of the BeeMonitor system at the Penn State Horticulture Research Farm, 2024. (A) Complete off-grid monitoring unit comprising the video recording module mounted on a tripod directed at the bee hotel, with the solar panel and battery enclosure positioned on the ground. (B) Nest tubes following collection at the end of the monitoring period, showing brood cells provisioned by *Osmia cornifrons*.

A population of approximately 300 cocoons of *Osmia cornifrons* (Watts Bees) were obtained during spring 2024 and maintained at 4 C in darkness until field use. Prior to their release, bees were maintained indoors at ambient temperature for approximately 48 hours to encourage adult emergence. Bees were then relocated to experimental sites along with a provided stacked, grooved wood board bee hotel (Fig 3B) at the start of the monitoring period (April 16 to May 16, 2024), enabling observation of foraging and nesting activity across a complete nesting cycle under natural environmental conditions.

The system recorded video continuously during daylight hours, generating high-resolution observations of bee activity across the monitoring period. From these recordings, we extracted foraging events (i.e., entry or exit of a bee from) and quantified foraging trips across temporal scales (e.g., min, hourly, daily).

Following the nesting period, reproductive output was quantified by counting the number of brood cells constructed in each nest tube (Figure 3B). These measurements provided an independent estimate of nesting activity, enabling comparison between observed foraging behavior and realized reproductive effort.

Environmental data were obtained from a nearby personal weather station (Weather Underground station KPABELLE49; 40.88° N, −77.83° W) at 5-minute intervals. Variables included temperature (°C), humidity (%), wind speed (m/s), atmospheric pressure (hPa), solar radiation (W/m²), and precipitation rate (mm/h), and were used to support analysis of environmental drivers of foraging behavior.

To quantify system hardware reliability in the field under adverse weather conditions, we calculated temporal coverage as the proportion of expected recording intervals for which video data were successfully captured and saved. The system was configured to record in 10-minute intervals between 09:00 and 19:00 each day, yielding 61 expected recording intervals per day. Coverage was computed as the fraction of these intervals with successfully recorded video.

### 2.4 Software pipeline evaluation

System performance was evaluated using a manually annotated dataset comprising 11 video segments (10 minutes each) selected from different days and times to represent a range of activity levels and environmental conditions. A trained observer reviewed each video to identify all foraging events, recording the event type (nest entry or exit), associated nest location, and timestamp. In total, 300 foraging events were annotated across the evaluation dataset. Pipeline performance was assessed at two levels. First, we evaluated the pipeline’s overall performance in accurately extracting foraging events from video. Second, we evaluate the generalization performance of the foraging-event machine-learning classifier in the activity-extraction module.

The overall performance was assessed using precision, recall, and F1 score. A predicted event was considered a true positive if a corresponding manually annotated event occurred at the same nest location within a ±2-second temporal window, accounting for small timing discrepancies between manual annotation and automated detection. Predictions without a matching annotation were counted as false positives, while annotated events without a corresponding prediction were counted as false negatives.

To evaluate the generalization of the machine learning event classifier across recording conditions, we performed leave-one-video-out (LOVO) cross-validation across all 11 annotated video segments. In each fold, the classifier was trained on trajectory data from 10 videos and evaluated on the held-out video; this procedure was repeated for all 11 folds. Performance was summarised using pooled precision, recall, and F1 score aggregated across all folds, as well as mean per-fold F1 and its standard deviation to characterize variability across recording conditions.

### 2.5 Evaluation of the environmental drivers of foraging and nesting activity

Foraging activity was quantified using foraging trips as the primary behavioral unit. Candidate trips were filtered to retain biologically plausible durations, excluding trips shorter than 0.5 min or longer than 30 min, consistent with observed foraging time scales for *Osmia cornifrons* (K. Y. Lee et al., 2010; McKinney & Park, 2012). Each foraging trip was assigned to the 5-minute interval corresponding to the nest exit event, which marked the initiation of the trip. Foraging trips were then aggregated at 5-minute intervals to match the temporal resolution of the weather data described in section 2.3, enabling direct alignment between behavioral activity and environmental conditions without temporal smoothing or interpolation.

Reproductive output was measured as the number of brood cells constructed per nest tube as described in section 2.3. To assess the relationship between foraging activity and reproductive effort, we conducted a correlation analysis of the total number of foraging trips and brood cell counts at the individual-nest level.

To quantify the influence of environmental conditions on foraging behavior, we modeled foraging trip initiation as a binary outcome, active or inactive per interval, using a Random Forest classifier. This approach captures nonlinear relationships and interactions between environmental drivers and behavioral responses. Partial dependence analysis was used to characterize the marginal effect of each environmental variable on foraging probability across its observed range. Model performance was evaluated using the area under the receiver operating characteristic curve (AUC) with five-fold time-series cross-validation to prevent data leakage across temporally autocorrelated observations.

## 3. Results

### 3.1 Field Deployment Hardware Reliability Evaluation

System reliability was evaluated using temporal coverage of recording and storage during the 2024 monitoring period (April 17–May 15). Recording coverage was consistently high across the monitoring period, with a mean (± SD) of 97.51% ± 10.28% and a range of 47.54–100.00% (Figure 4A). Most days showed near-complete coverage throughout daylight hours, with only isolated gaps due to brief system interruptions.

**Figure 4.**
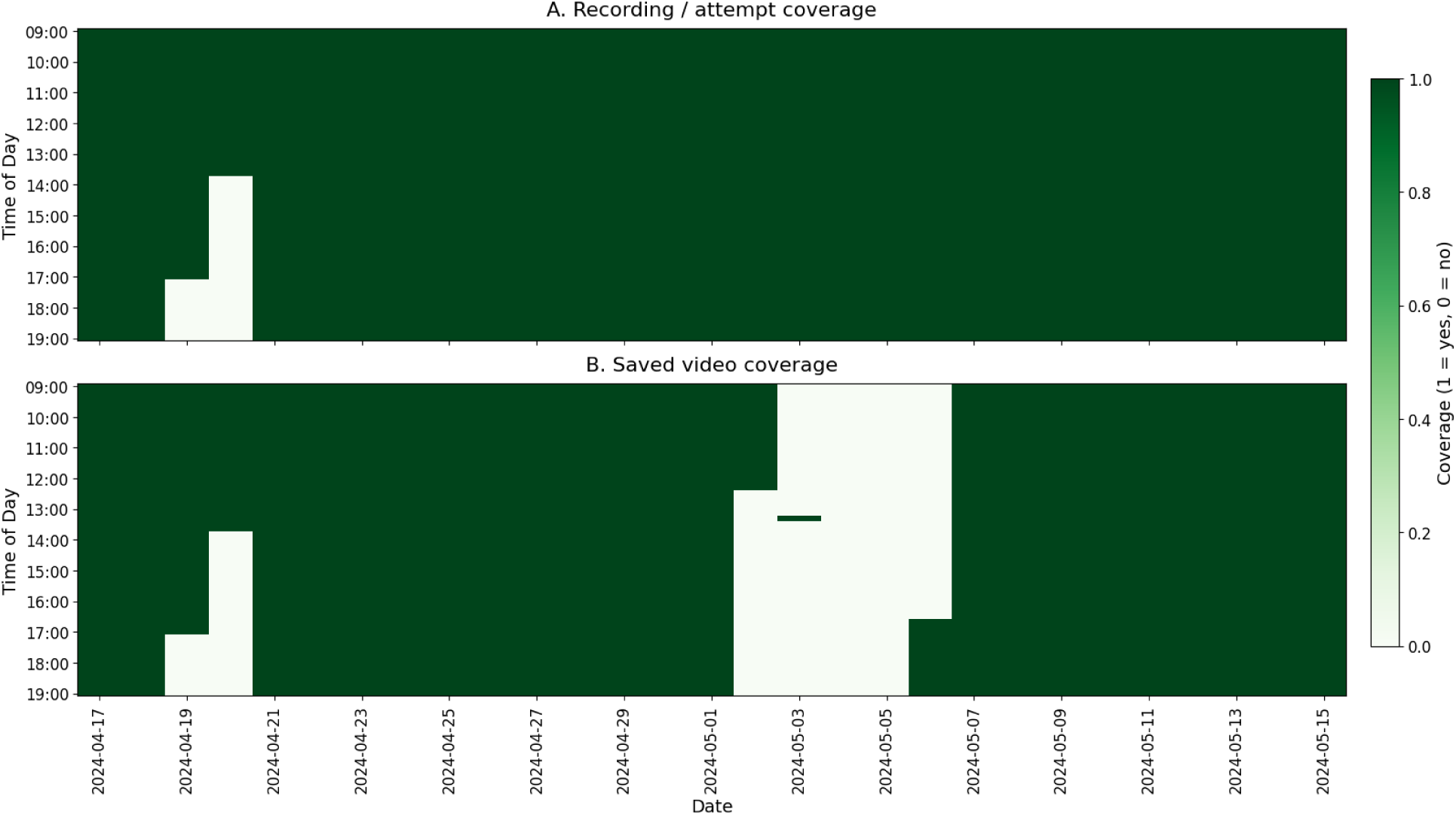
Temporal coverage of the BeeMonitor recording system, April 17 – May 15, 2024. Each cell represents a 10-minute recording interval between 09:00 and 19:00, with green indicating a successfully recorded interval and white indicating a gap. (A) Recording attempt coverage reflects the intervals during which the system logged a recording event. Coverage was consistently high throughout the monitoring period, with isolated gaps on April 19–20 due to intermittent system interruptions during early deployment. (B) Saved video coverage, reflecting intervals for which video data were successfully written to storage. A pronounced gap between approximately May 2–6 corresponds to a period of storage limitation during which the system continued operating, but recordings were not successfully saved.

Successfully saved video coverage was lower and more variable, with a mean of 82.36% ± 34.73% and a range of 0.00–100.00% (Figure 4B). A pronounced gap between approximately May 2 and May 6 corresponded to a period of storage limitation, during which recordings were not successfully saved despite continued system operation. Smaller gaps on April 19–20 were associated with intermittent system interruptions during early deployment. Outside these periods, saved video coverage was consistently high, demonstrating reliable system operation under field conditions.

### 3.2 Video Processing Software Pipeline Evaluation

The BeeMonitor computer vision software pipeline was evaluated using 11 manually annotated video segments spanning multiple days and times of day. The pipeline achieved a precision of 91.3%, a recall of 87.3%, and an F1 score of 0.893, correctly identifying 262 of 300 annotated events, with 25 false positives and 38 false negatives (Figure 5). Entry events were detected with slightly higher performance (precision 94.4%, recall 91.3%, F1 0.928) than exit events (precision 88.1%, recall 83.4%, F1 0.857).

**Figure 5.**
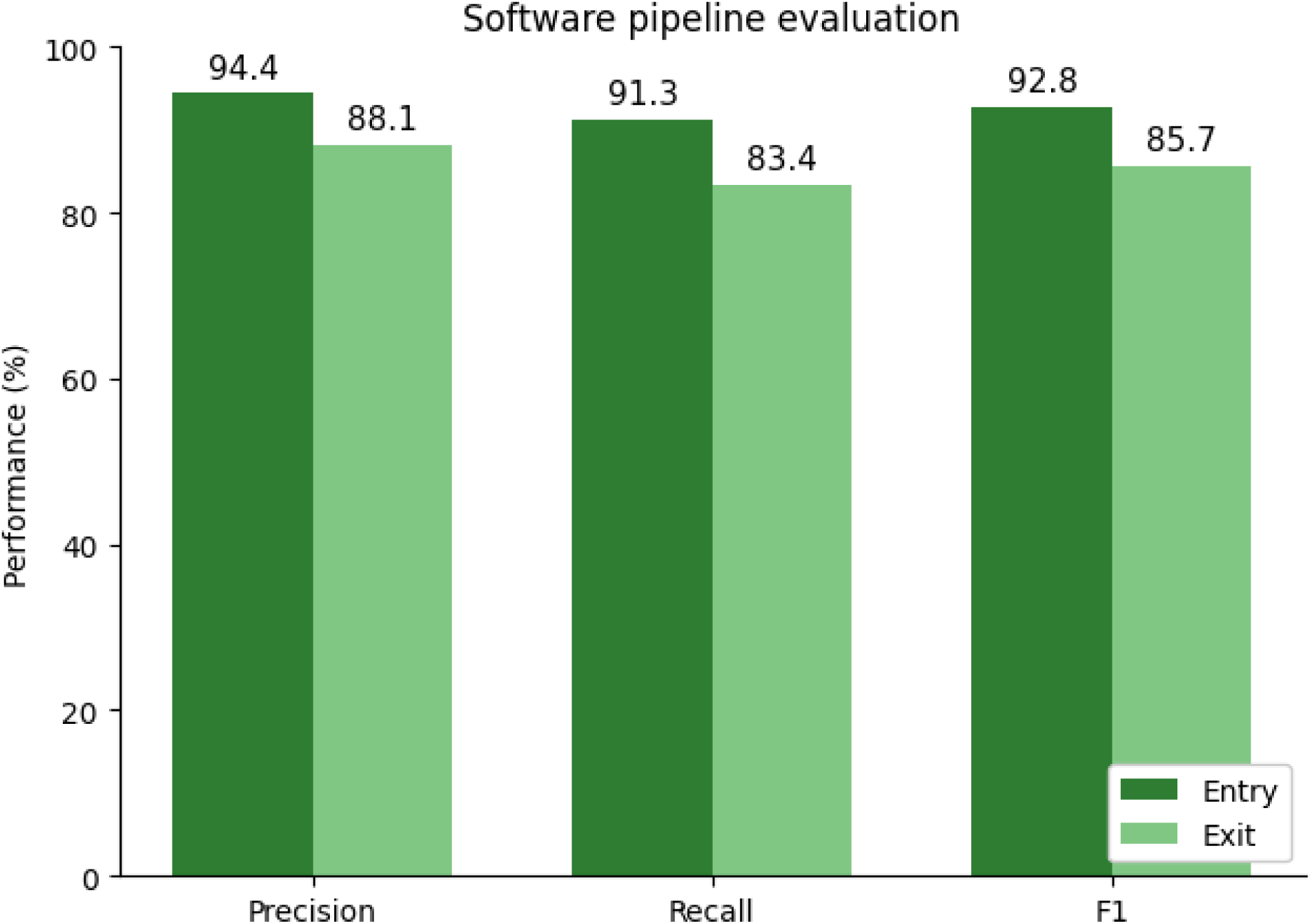
Software pipeline evaluation performance by event type. Precision, recall, and F1 score are shown separately for nest entry and exit events, evaluated against 300 manually annotated foraging events across 11 video segments using a ±2 s matching window. Entry events were detected with consistently higher performance than exit events across all three metrics, reflecting the more distinctive trajectory characteristics of bees approaching and entering nest tubes compared to departing bees whose trajectories begin inside the nest.

To evaluate the generalization of the machine learning event classifier across recording conditions, leave-one-video-out (LOVO) cross-validation was performed across all 11 video segments. The classifier achieved a pooled precision of 90.4%, recall of 90.8%, and F1 score of 0.906, with a mean per-fold F1 of 0.904 ± 0.045 and individual video F1 scores ranging from 0.811 to 0.971 (Table 1). Performance was consistent across videos recorded under varying activity levels, times of day, and weather conditions, demonstrating robust generalization of the classifier to unseen recording conditions.

**Table 1.** Leave-one-video-out (LOVO) cross-validation results for the machine learning event classifier. Each row represents one held-out video fold, identified by recording date and start time. Precision, recall, and F1 score are reported for each fold, with the pooled row showing aggregate metrics across all 11 folds. Evaluation used a ±2 matching window at the same nest location and event type.

| Video | Precision | Recall | F1 |
| --- | --- | --- | --- |
| 2024-04-30 09:00 | 1.000 | 0.889 | 0.941 |
| 2024-04-30 09:10 | 0.923 | 0.857 | 0.889 |
| 2024-04-30 09:20 | 0.875 | 1.000 | 0.933 |
| 2024-04-30 09:30 | 1.000 | 0.882 | 0.938 |
| 2024-04-30 09:40 | 0.750 | 1.000 | 0.857 |
| 2024-05-08 15:00 | 0.938 | 0.714 | 0.811 |
| 2024-05-08 15:30 | 0.955 | 0.913 | 0.933 |
| 2024-05-08 15:50 | 0.840 | 0.955 | 0.894 |
| 2024-05-23 12:00 | 0.917 | 0.917 | 0.917 |
| 2024-05-23 12:40 | 0.962 | 0.981 | 0.971 |
| 2024-05-23 18:20 | 0.823 | 0.895 | 0.857 |
| Pooled | 0.904 | 0.908 | 0.906 |

### 3.3 Foraging activity and reproductive output

A strong positive relationship was observed between foraging activity and reproductive output at the nest tube level. Spatial patterns across the bee hotel showed that nest tubes with higher foraging activity also tended to have more brood cells (Figure 6). Across all monitored nest tubes, the number of brood cells per nest increased with the total number of foraging trips detected by BeeMonitor (R² = 0.849, p < 0.001, n = 19 nest tubes; Figure 7). The estimated slope of 0.07 brood cells per foraging trip indicates that approximately 14 foraging trips were required to provision a single brood cell.

**Figure 6.**
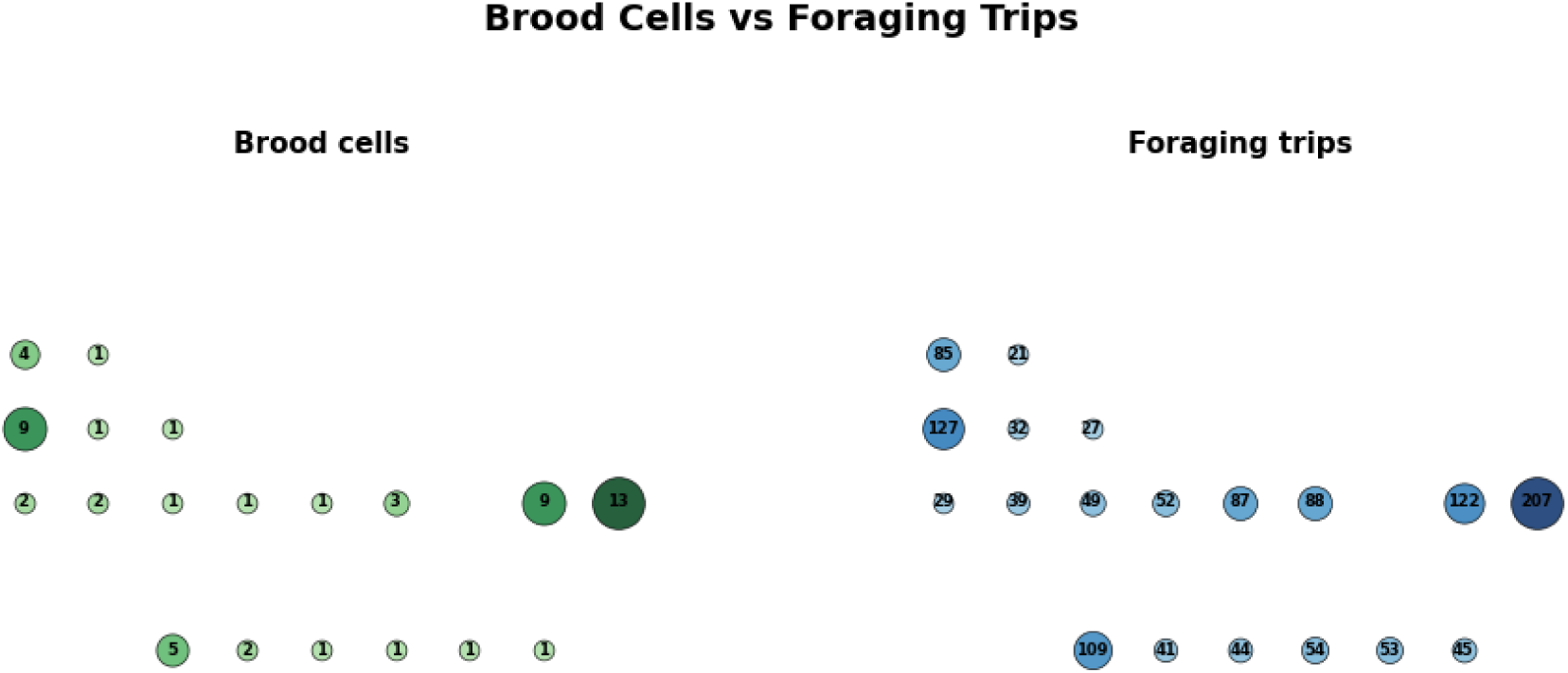
Spatial distribution of brood cells and foraging trips across the bee hotel. The bee hotel had 10 columns and 6 rows creating 60 nest tubes. Each circle represents a nest tube, with circle size proportional to the count of brood cells or total foraging trips for the nest tube.

**Figure 7.**
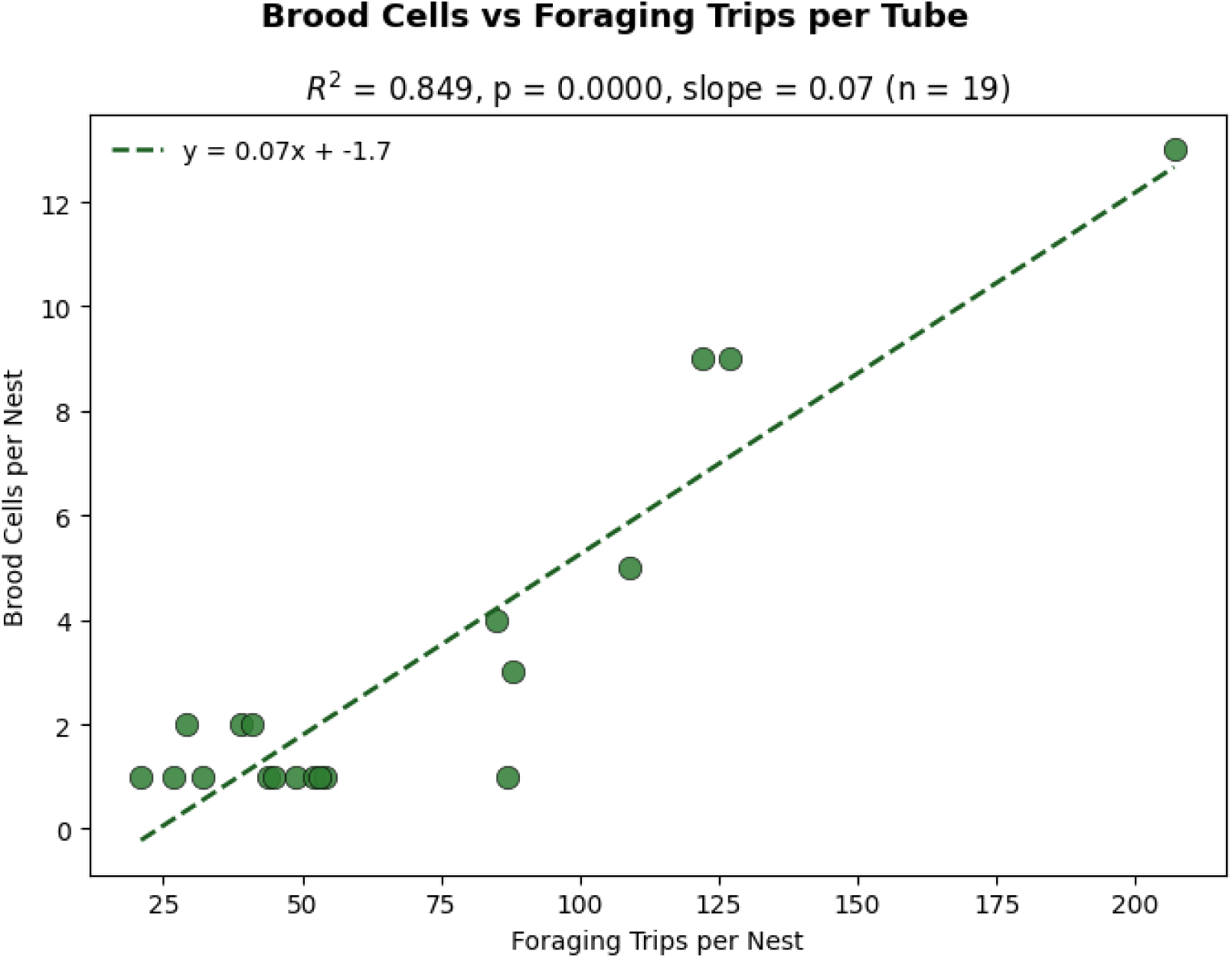
Relationship between foraging activity and reproductive output at the nest tube level. Each point represents one nest tube (n = 19). The dashed line shows the fitted linear regression (y = 0.07x − 1.7). The strong positive relationship (R² = 0.849, p < 0.001) indicates that approximately 14 foraging trips were required to provision a single brood cell.

### 3.4 Environmental drivers of foraging activity

The Random Forest classifier achieved good predictive performance (mean AUC = 0.820 ± 0.062; n = 4,859 five-minute intervals), indicating that environmental variables captured substantial variation in foraging activity across the monitoring period. Feature importance analysis identified solar radiation as the dominant predictor of foraging activity (36.7%), followed by temperature (26.0%), humidity (18.7%), atmospheric pressure (12.2%), wind speed (6.4%), and precipitation rate (0.1%; Figure 8).

**Figure 8.**
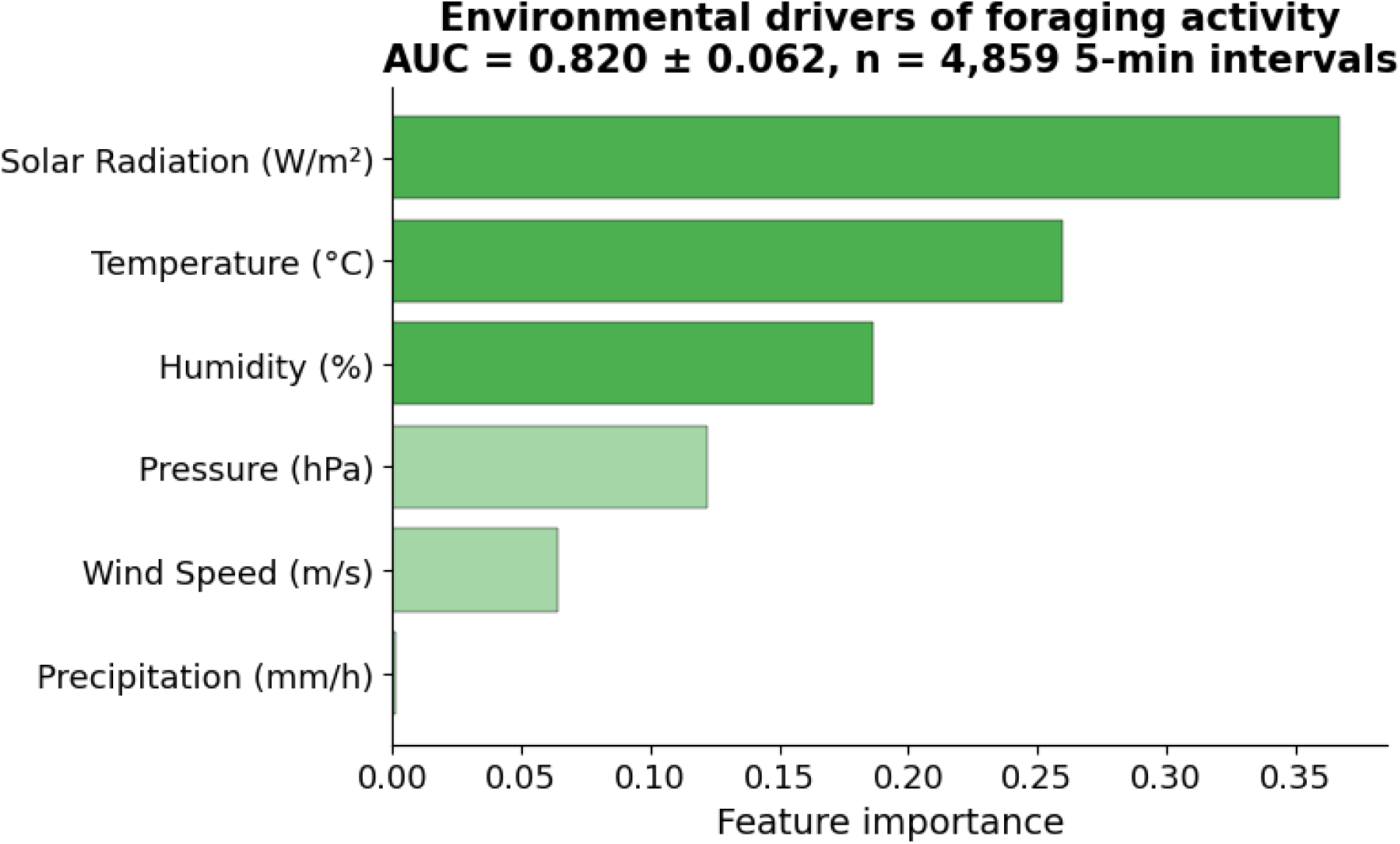
Relative importance of environmental variables for predicting foraging activity in *Osmia cornifrons*. Feature importance scores are derived from a Random Forest classifier trained on 4,859 five-minute intervals (AUC = 0.820 ± 0.062). Darker bars indicate variables with above-median importance. Solar radiation was the dominant predictor, followed by temperature and humidity.

## 4. Discussion

Quantifying the foraging behavior of cavity-nesting solitary bees at fine spatiotemporal scales has remained a persistent challenge for pollinator research, management, and conservation, largely due to the labor-intensive nature of manual observation and its inherent limitations in spatial and temporal coverage and resolution. BeeMonitor addresses this gap by providing an automated, continuous monitoring system that extracts discrete nest-tube-level foraging activity from video recordings without requiring manual video processing. The results presented here provide evidence that the BeeMonitor can reliably capture foraging behavior throughout a complete nesting season.

### 4.1 Hardware reliability and field viability

The recording system achieved a mean recording coverage of 97.51% across the monitoring period, demonstrating that the hardware is sufficiently reliable for deployment in real field conditions. The pronounced gap in saved video coverage between May 2-6, attributable to storage limitations rather than system failure, highlights an important practical consideration for long-term deployments. Outside this period, saved video coverage was consistently high, confirming that the system can sustain continuous daylight monitoring over a complete nesting season.

### 4.2 Pipeline performance and comparison to existing systems

The BeeMonitor software pipeline achieved an overall precision of 91.3% and recall of 87.3% (F1 = 0.893), with nest tube entry activity detected more accurately than exit activity. This performance gap reflects the inherently challenging nature of exit detection. Bees depart rapidly from the nest tube entrance, producing shorter observable trajectories that make activity detection difficult. Increasing camera frame rate per second could address this challenge in the future. Compared to existing approaches, the BeeMonitor software bridges an important gap. Systems requiring individual marking or identification tags, such as Bee Tracker (Knauer et al., 2022), achieves high precision of 96% for detecting tagged *Osmia bicornis* foraging activity at a bee hotel. However, Bee Tracker evaluation does not report recall of system and requires challenging labor to tag bees, which can also affect bees’ natural foraging and nesting behavior. On the other hand, other video monitoring systems for solitary bees that do not require bee tags achieves low precision of 78% (Wittmann et al., 2024). BeeMonitor achieves a precision of 91.3% and a recall of 87.3% on untagged *Osmia cornifrons* under natural field conditions, thus achieving high accuracy without labor intensive manual review of videos or tagging of bees.

### 4.3 Ecological validation

The strong positive relationship between BeeMonitor-detected foraging trips (i.e., consecutive exit and entry into a nest tube) and brood cell counts (R² = 0.849, p < 0.001) provides robust validation that the system captures biologically meaningful variation of nesting activity. Our estimated rate of approximately 14 foraging trips per brood cell is lower than the 31.3 trips per cell reported by McKinney & Park (2012). This difference may reflect variation in local resource availability and landscape context between study sites. Foraging trip duration and frequency can be sensitive to landscape-level resource abundance (Hemberger & Gratton, 2018; Westphal et al., 2006). Despite this difference, the consistent positive relationship across 19 nest tubes provide strong evidence that BeeMonitor-derived foraging metrics are a reliable proxy for reproductive effort at the nest level.

### 4.4 Environmental drivers of foraging activity

Foraging activity of *Osmia cornifrons* is strongly influenced by solar radiation and temperature. Together, these two factors explained 62.7% of the variation in *Osmia cornifrons* egress from a bee hotel to forage. The results are consistent with other modeling studies, which have found that 78% of honey bee egress from a hive can be explained by solar radiation and temperature (Clarke & Robert, 2018). Bees depend on the sun for navigation and solar radiation contributes significantly to bee body temperature, which may explain why solar radiation emerged as a dominant predictor for *Osmia cornifrons* foraging activity(Johnson et al., 2025; Waiker et al., 2019). Further, the limited influence of wind speed and precipitation as a driver of foraging activity upholds the widely held but largely anecdotal view that *Osmia* spp. foraging habits are generally unaffected by light wind or rain (Vicens & Bosch, 2000).

### 4.5 Limitations and future directions

This study was conducted at a single site over a single nesting season, which limits the generalisability of the ecological findings. Replication across multiple sites, years, and with different cavity-nesting bee species would establish the robustness of the environmental relationships identified here and enable species-specific comparisons of foraging responses to environmental change (Hamblin et al., 2017; E. Lee et al., 2018). The storage limitation observed in early May highlights the need for automated cloud-based data transfer in future deployments, which would enable truly continuous remote monitoring without dependence on physical site visits (Knauer et al., 2022; Wittmann et al., 2024). Future versions of BeeMonitor will also incorporate onboard environmental sensing directly into the hardware unit, eliminating dependence on nearby weather stations. Finally, a taxonomic and a tagged-bee identification system will be incorporated into BeeMonitor to track individual bees.

## Author Contributions

EIA, SS, NB, and CMG conceived BeeMonitor. EIA, SS, and NB developed BeeMonitor. EIA and SS deployed the BeeMonitor. EIA, SS, NB, and CMG wrote and revised the manuscript. NB and CMG supervised the work. All authors approved the final version.

## Acknowledgements

We want to thank everyone in the Christina Grozinger lab for helpful and thoughtful feedback. Special thanks to Jaya Sravanthi Mokkapati for her assistance with processing nesting data from the field. We would also like to thank Orion Pizzini for his assistance with setting up the bee hotel and camera system in the field. The authors of this work recognize the Penn State Institute for Computational and Data Sciences (RRID:SCR_025154) for providing access to computational research infrastructure within the Roar Core Facility (RRID: SCR_026424).

## Funding

Funding for EA was provided by the Interdisciplinary Studies in Entomology, Computer Science and Technology Network (INSECT NET) supported by the National Science Foundation’s Research Traineeship Program (Grozinger and Boyle, Grant 2243979). Additional funding was provided by the USDA National Institute of Food and Agriculture, Hatch and Smith-Lever Appropriations under Projects PEN04943 and PEN08801, and by Penn State’s Joan Luerssen Faculty Enhancement Fund.

## Conflict of interest

The authors declare no competing interests.

## Data and code availability

The BeeMonitor software is accessible at https://mqnafc3ejc.us-east-1.awsapprunner.com/accounts/login/?next<u>=/</u> . Source code for software and 3D stl files can be found on the official GitHub repository here https://github.com/Team-Insect-Net/BeeMonitor. Validation datasets and code are available on Scholars Sphere here https://scholarsphere.psu.edu/resources/55f1f34b-959f-4c60-8dd3-9b33fb09357f.

## Notes

https://scholarsphere.psu.edu/resources/55f1f34b-959f-4c60-8dd3-9b33fb09357f

### Competing Interest Statement

The authors have declared no competing interest.

